# SUMOylation of SARS-CoV-2 Nucleocapsid protein enhances its interaction affinity and plays a critical role for its nuclear translocation

**DOI:** 10.1101/2022.05.31.494262

**Authors:** Vipul Madahar, Victor G. J. Rodgers, Jiayu Liao

## Abstract

Viruses, such as SARS-CoV-2, infect hosts and take advantages of host cellular machinery for their genome replication and new virion production. Identification and elucidation of host pathways for viral infection are critical for understanding the viral life cycle and novel therapeutics development. SARS-CoV-2 N protein is critical for viral RNA(vRNA) genome packaging in new virion formation, Here, we report that identification of SUMOylation sites of SARS-CoV-2 N protein and role of SUMO modification in N protein interaction affinity with itself using our qFRET/MS coupled method. We found, for the first time, that the SUMO modification of N protein can significantly increase its interaction affinity with itself and may support its oligomer formation. One of the identified Lys residues, K65 was critical for N protein translocation to nucleus, where the vRNA replication and packaging take place. The *in vitro* assessment of the affinity of N protein to N protein with SUMO mutants provides insight of the oligomerized N protein formation after SUMO modification. These results suggest that the host human SUMOylation pathway may be very critical for N protein functions in viral replication. The host SUMOylation pathway may be a critical host factor for the SARS-CoV-2 virus life cycle. Identification and inhibition of critical host SUMOyaltion could provide a novel strategy for future anti-viral therapeutics development, such as SARS-CoV-2 and other viruses.

**Importance:** The SARS-CoV-2 virus N protein plays a critical role critical for viral RNA(vRNA) genome packaging in host cell nucleus for new virion formation. Therefore, deciphering the molecular mechanisms modulating N activity could be a strategy to identify potential targets amenable to therapeutics. Here, we identify a comprehensive SUMOylation sites of N proteins using an *in vitro* reconstitute SUMOyaltion assay containing SUMO E1 activating enzyme, E2 conjugating enzyme, and E3 ligase. We find that SUMOylation modification of N protein can significantly enhance it interaction affinity with itself, indicating an increased oligomerization capability, which is critical for N protein activity for vRNA genome packaging. In addition, we find one of SUMOylation sites of N protein is critical for its nucleus translocation, which is a critical for viral genome packaging. The SUMOylation modification may represent novel potential approach to design new antivirals with the ability to modulate SARS-CoV-2 virus replication.

## Introduction

The severe acute respiratory syndrome coronavirus 2 (SARS-CoV-2) is a global pandemic responsible for the upper respiratory disease Coronavirus Disease 2019 (COVID-19). The rapid development of variants from the original strain with onset of functional mutations highlights to the need to discover the pathogenesis and potential new strategy of anti-viral therapeutics development. The SARS-CoV-2 viral particle is composed of the positive sense single strand RNA (∼30 kb) genome, which encode 29 proteins, packed around nucleocapsid (N) proteins. The compacted RNA nucleocapsid complex is enveloped in a lipid membrane with embedded membrane proteins (M) and envelope protein (E)(1). The glycoprotein spike protein (S) exposed outside on the viral particle surface is found to have high affinity for human angiotensin-converting enzyme 2 (hACE2)(2).

Recent studies have demonstrated that the post translation modification (PTM) glycosylation plays an essential role in S and hACE2 interaction(3). Once bound, the human cell surface protein transmembrane protease serine 2 (TMPRSS2) cleaves the spike protein which results in the cleaved spike protein binding to the host cell surface(2). The S protein unravels and merges the two lipid membranes together and an endocytosis mechanism delivers the particle into the cell(4). Recent PTM studies on the SARS-CoV-2 N protein have found potential phosphorylation on serine 197 and threonine 205(5). The study observed SARS-CoV-2 N protein modulation of RNA binding with mutations at 197 and 205 in the S/R rich region. The phosphorylation modification is inherent to the proteins function and demonstrates the dependence of virus progression on host PTMs(6, 7). Furthermore, an investigation in ubiquitination, screened in-cell modified N protein for ubiquitination and observed lysine 169, 374, and 388 to be ubiquitin modification, but no follow up functional study(8).

SARS-CoV N protein is critical for binding vRNA for *ribonucleoprotein (RNP) assembly* and interact with M and E proteins to help viral envelop formation and viral particle assembly(9, 10). Cryo-electron tomography (CryoET) studies have elucidated the complex overall structure of N protein bound to vRNA and N protein bound to M protein. Recent investigations concluded the oligomerized N protein compacts RNA into a structure that forms a phase-separated condensate within the viral particle that binds to M protein(1, 6, 7, 11). They observed the average diameter of a viral particle to be 80 nm and contained 30-35 vRNPs. The group predicted each vRNP is 15 nm in diameter and holds 12 copies of N protein wrap and coated in RNA, creating a condensate that encapsulates the genomic vRNA and interacts with M protein(6, 12). The sub-domains that enable protein-protein complexes of the N protein oligomers to form dimers and tetramers, enables the complex formation of vRNP and ultimately enclosure within a viron.

The SARS-CoV N protein and SARS-CoV-2 N proteins have homology across their sequences and demonstrated to have similar domains and function(13). The N protein identified domains N-terminal domain (NTD from 1-50 aa) is followed by the RNA binding domain (RBD from 51-174 aa), the dimerization domain (247-364 aa), and the C-terminal domain (CTD from 365-419aa)(12). The CTD reported to interact with M protein within the viral envelope and supports the vRNA super structure(6). Between the RBD and the dimerization domain is a liker region (174-245 aa), this region is a serine and arginine rich region on the protein that is reported to be phosphorylated(5).

Investigations of the intersection of N protein interaction network with host proteome have expanded the functional attributes of N protein beyond the vRNP complex. The anti-viral response of host cells is reported to be inhibited by N protein interaction with host immune response protein, signal transducer and activator of transcription (STAT1 & 2). N protein mediate inhibition of interferon antiviral responses by sequestering activated STAT proteins within the cytosol and disrupting the IFN signaling from progressing(14). Furthermore, N protein is reported to depend on PTMs from host proteome for its functional RNA binding properties. The PTM SUMOylation on SARS-CoV N protein modulate its subcellular localization in cell(15). A significant impact that was alluded to in the 2005 study by Li et al is the modulation of oligomerization of N protein with the SUMO modification(15). The evidence provided in the study was a western blot that observed the decrease of crosslinked N proteins. However, it has been demonstrated that N proteins can form super structures of dimers and tetramer which ultimately form condensate, and the oligomerization of N protein is described to be a critical factor in viral genome packaging (6, 11)..

The SUMO modification of a target protein increases the target protein affinity to other cellular processes and extends the target proteins non-covalent interaction network range to proteins with affinity to SUMOylated proteins(16, 17). In this work we demonstrate an in-vitro qFRET assay for the SUMOylation of N protein coupled with mass spectrometry to identify the sites of SUMO modified lysine. One of the modified lysine sites, Lys65, when mutated to Arg, restricted the N protein in the cytosol, suggesting the SUMOylation may play a critical role in nuclear translocation of the N protein. In addition, we found SUMOylated N protein has much higher affinity to interact to each other than the un-SUMOylated N proteins. These results suggest that host SUMOylation may play important roles in SARS-CoV-2 life cycle and could be one potential target for future therapeutic development.

## Materials and Methods

### Expression and Purification of SUMOylation Enzymes and SARS-CoV-2 N protein

The in-vitro qFRET SUMOylation reaction is completed with the E1, E2, and E3 enzymes in the SUMOylation cascade. The E1 activation enzyme complex, UBA2 and AOS1, E2 conjugating enzyme UBC9, and E3 ligase PIAS1 were all cloned into pET28B vector for expression in BL21(DE3) cells. The FRET pairs CyPet and YPet are N-terminal tagged to SUMO1 and the substrates respectively and cloned into pET28B for expression in BL21(DE3). Each BL21(DE3) cell line with individual proteins were inoculated at 1:100 and grown to 0.4 OD at 600 nm at 37 °C, and then induced at overnight at 22°C with 0.25 mM IPTG. The cells were lysed (lysis buffer, 20 mM Tris-HCl (pH 7.5), 0.5 M NaCl, 5 mM Imidazole), by sonication and centrifuged at 35,000 x g. The soluble fraction was purified by 6XHis tag to NiNTA beads affinity chromatography through a gravity column. The bound proteins were washed with, buffer 1 (20 mM Tris-HCl (pH 7.5), 0.3 M NaCl), buffer 2 (20 mM Tris-HCl (pH 7.5), 1.5 M NaCl, and 0.5% Triton X-100), and buffer 3 (20 mM Tris-HCl pH 7.5, 0.5 M NaCl, and 10 mM Imidazole). The proteins eluted using the following buffer, (20 mM Tris-HCl, 300 mM NaCl, and 450 mM Imidazole) and dialyzed in 20 mM Tris-HCl (pH 7.5), 50 mM NaCl, and 1 mM DTT.

### In-vitro SUMOylation Assay Setup

The *in-vitro* SUMOylation assay is completed with 6xHisCyPet-SUMO1 500 nM, 6xHisYPet-N protein wildtype 2000 nM, E1 hetro-dimer AOS1/UBA2 at 100 nM, E2 conjugating enzyme UBC9 200 nM, E3 ligase PIAS1 250 nM, and in SUMOylation buffer (20 mM Tris-HCl (pH 7.5) 50 mM NaCl, 4 mM MgCl, 1 mM DTT). Functional controls are put in place for observing non-specific interaction, by a negative control reaction without 2 mM adenosine triphosphate (ATP). Each reaction was incubated at 37°C for 60 minutes and measured in a 384 well microplate (Grenier 384 M6811). The FRET wavelength, E_mTotal_, are 414 nm excitation and 530 nm emission, Fl_DD_, 414 nm excitation and 475 nm emission, and Fl_(18)A_, 475 nm excitation and 530 nm emission. The quantitative E_mFRET_ parameters, α of 0.34 +/- 0.003, and β 0.003 +/- 0.001 variables are determined using the formulation outlined in previous work from Yang et al(18). Equation 1 provides the calculation of E_mFRET_ that quantifies the FRET signal by substracting free donor and acceptor emissions from the total fluorescence emission.

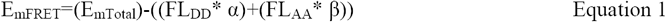

The specificity of SUMO protein to the SUMOylation target has the potential to yield a false positive FRET response. Thus, functional controls of reactions without ATP are implemented in parallel to observe differences between ATP and no ATP. Samples from each reaction of -ATP/- E3/+ATP+E3 was also immune blotted with anti SUMO1.

### Mass spectrometry analysis to determine SUMO modified Lysine on N protein

The *in-vitro* SUMOylation reactions the substate, YPet-SARS-CoV-2 Nucleocapsid protein is added at 3000 nM, and CyPet tagged SUMO1 protein are added at 1000 nM. Activating Enzyme Complex 1 (E1) is at 100 nM, and Conjugating Enzyme 2 (E2) at 100 nM, E3 ligase at 500 nM in SUMOylation buffer (20 mM Tris-HCl (pH 7.5) 50 mM NaCl, 4 mM MgCl, 1 mM DTT) and 2 mM ATP. The reactions were completed at 37° C for 4 hours. The in-solution proteolytic digestions were performed with Pierce™ Glu-C Protease. Samples were digested at 1:100 ratio for sample to enzyme ratio and ran overnight (16 hours) at 37°C. Each completed digestion was acidified to a final concentration of 0.1% v/v TFA, and speed vacuumed to dry product, and then reconstituted to 0.1% v/v TFA readied for MS loading.

### LTQ Orbitrap Xl Loading and Run

Samples consisted of approximately 1000 nM of in-solution digested product. From each proteolytic, enzyme digestion liquid chromatography was performed on a Thermo nLC1200 (ThermoFisher™) in single-pump trapping mode with a Thermo PepMap RSLC C18 EASY-spray column (2 μm, 100 Å, 75 μm x 25 cm) and a Pepmap C18 trap column (3 μm, 100 Å, 75 μm x 20 mm). Solvents used were A: water with 0.1% formic acid and B: 80% acetonitrile with 0.1% formic acid. Samples were separated at 300 nL/min with a 250-minute gradient starting at 3% B increasing to 30% B from 1 to 231 minutes, then to 85% B at 241 minutes, holding for 10 minutes.

Mass spectrometry data was acquired on a Thermo Orbitrap Fusion (ThermoFisher™) in data-dependent mode. A full scan was conducted using 60k resolution in the Orbitrap in positive mode. Precursors for MS^2^ were filtered by monoisotopic peak determination for peptides, intensity threshold 5.0 × 10^3^, charge state 2-7, and 60 second dynamic exclusion after 1 analysis with a mass tolerance of 10 ppm. Higher-energy C-trap dissociation (HCD) spectra were collected in ion trap MS^2^ at 35% energy and isolation window 1.6 m/z.

### Bioinformatic Analysis of MS Data

The LTQ-orbitrap XL (.raw) raw data was analyzed on Thermofisher Proteome AnalyzerTM. The complete amino acid sequence of each protein was provided as a reference for analysis. The expected SUMOylated lysines proteolytic products are tabulated (Table 1)ated and matched to mass over charge spectrums within the Thermofisher Proteome AnalyzerTMsearched for usingng both software suites. Precursor ion peptide tolerances were set at 5 ppm, and MS/MS peptide tolerances were set at 1 Dalton.

**Table 1:**
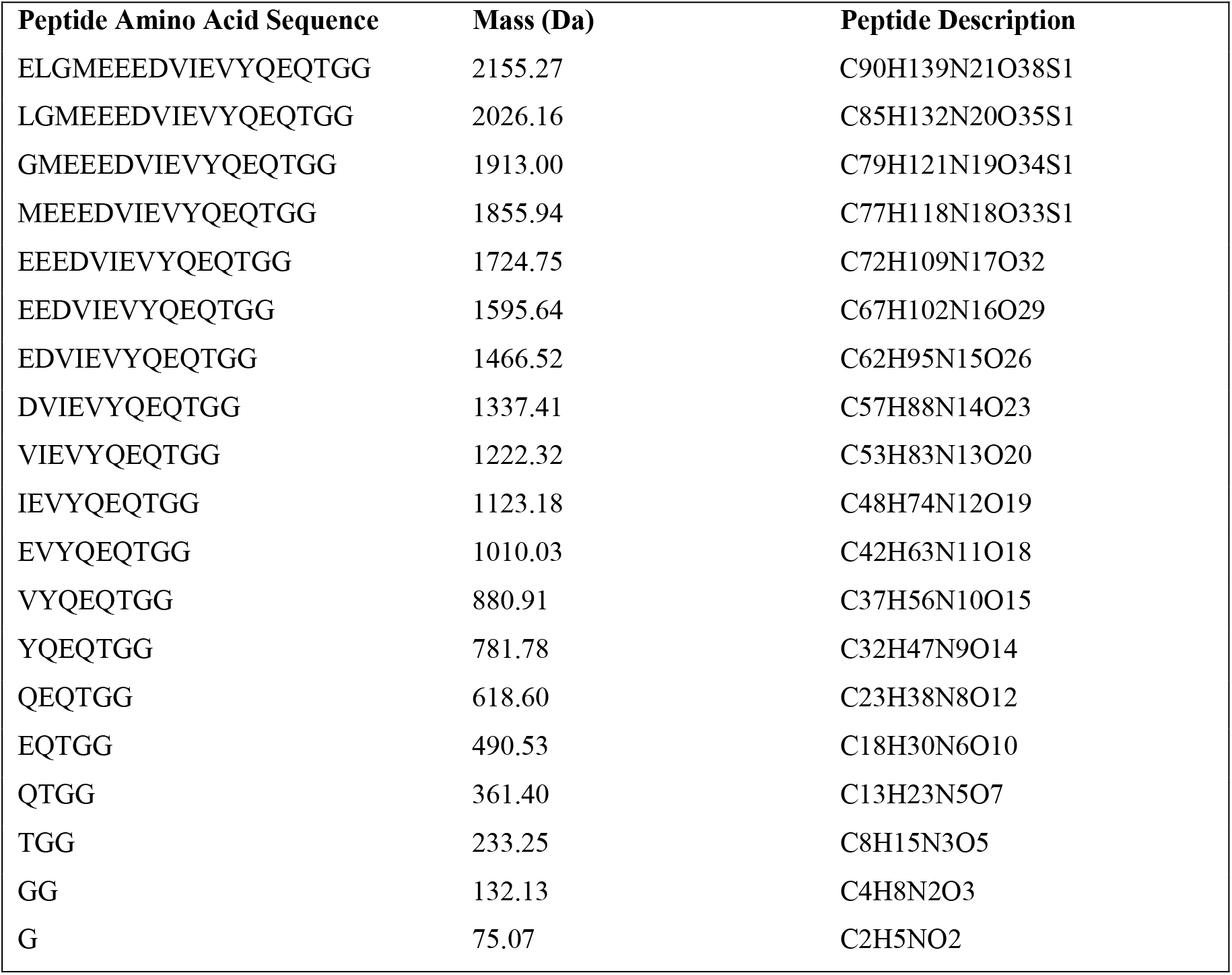
SUMO1 proteolytic peptides for identification of lysine modification.

### Construction and Design of N Protein Lysine to Arginine Mutants

The mass spectrometry results provided a total of four lysine residues that were SUMOylated. There are a total of 31 lysine residues on SARS-CoV-2 N protein, Lysine 61, 65, 347, and 355 were found to be SUMOylated in the *in vitro* reaction. The mutants DNA templates were constructed through PCR, with point mutations at the lysine to arginine coding sequences. The primers for the mutations were listed in Table 2. Final Gibson reaction of bacterial expression pET28B vector SALI and NOTI and for mammalian expression pCDNA3.1-FLAGtag-Nprotein- YPet were created. Tabulated PCR primers are shown in Table 3 for pcDNA3.1.

**Table 2:**
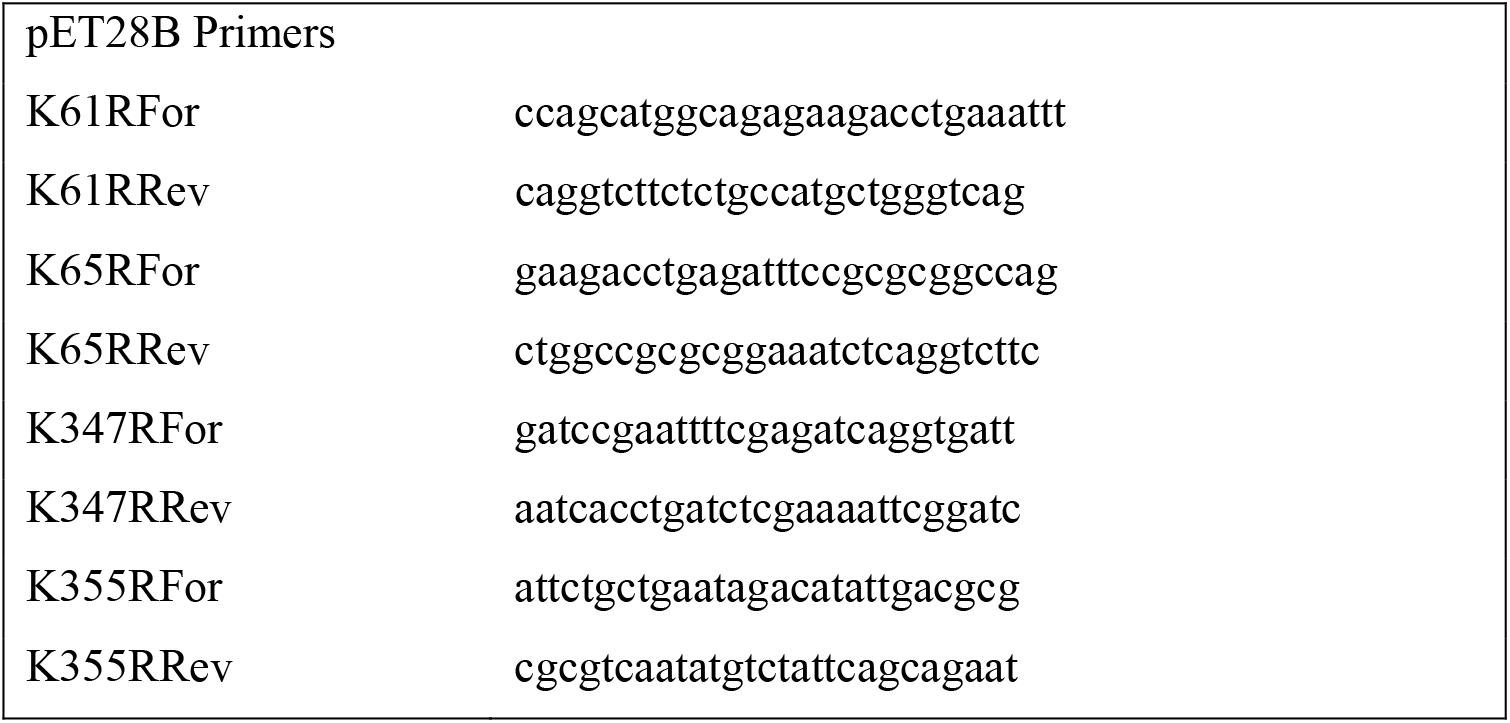
Primers listed for constructing N Protein Mutants in E. coli.

**Table 3:**
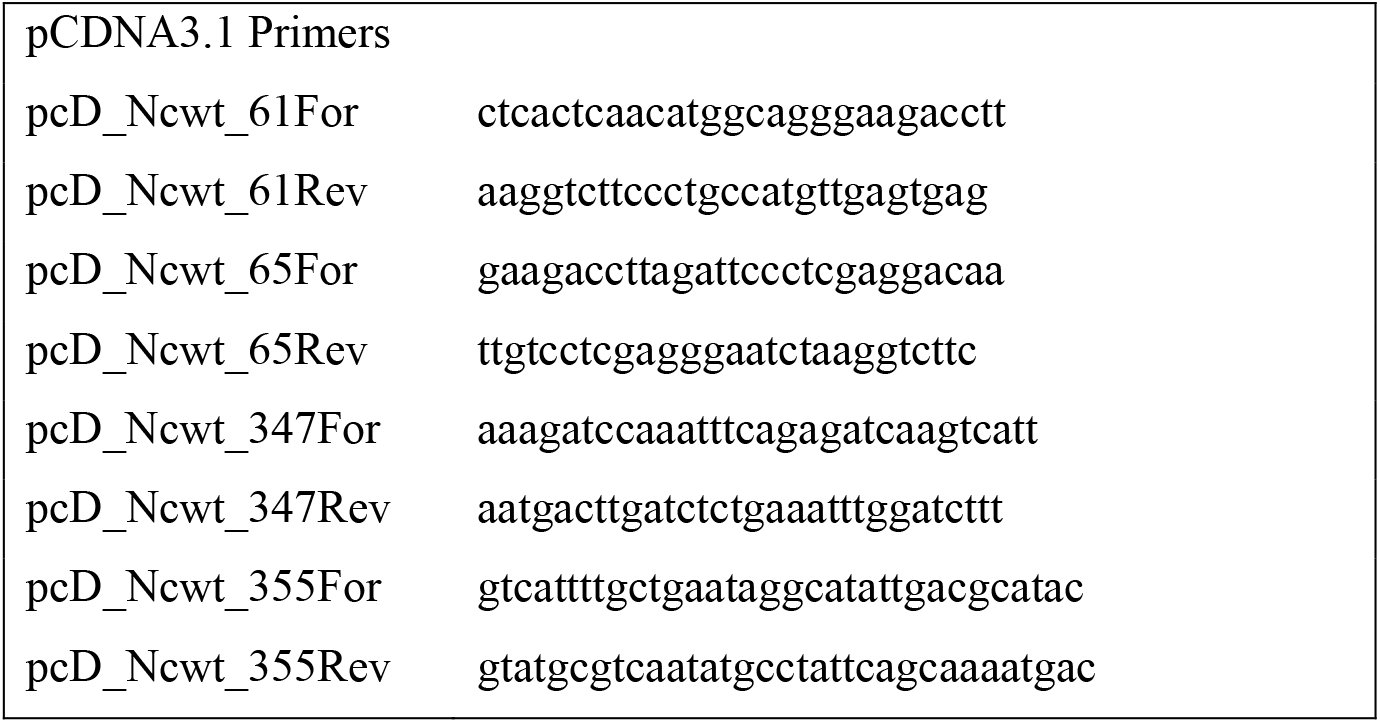
List of primers for mutations on N protein in HUH7 cells.

### *In-Vitro* SUMOylation with qFRET Reporter for N Protein Mutants

The in-vitro SUMOylation assay of SARS-CoV-2 N protein mutants is an initial screening to determine impact of lysine sites on SUMOylation. The assay is setup at the same concentration as the optimized conditions, 6xHisCyPet-SUMO1 500 nM, 6xHisYPet-N Protein wildtype and mutants 2000 nM, E1 hetro-dimer AOS1/UBA2 at 100 nM, E2 conjugating enzyme UBC9 200 nM, E3 ligase PIAS1 250 nM, and SUMOylation buffer of 20 mM Tris-HCl (pH 7.5) 50 mM NaCl, 4 mM MgCl, 1 mM DTT. Functional controls are put in place for non-specific interaction, by a negative control reaction without ATP, and to observe a significant boost in FRET a control reaction without E3 ligase. Each reaction was incubated at 37°C for 60 minutes. Following Equation 1 we measure the three fluorescence emissions, E_mTotal_, FL_DD_, and FL_AA_. The measurements are taken on Molecular Devices Spectra M3™, with “Endpoint” settings, with PMT at constant gain set to “Low”. The post analysis was completed on GraphpadPrism7™, One-way ANOVA with post HOC Tukey Test was done with minus ATP as the control group.

### Cellular Translocation of N protein

Immunostaining of N protein is used to investigate the dependency of SUMOylation of N protein on translocation between cytosol and nucleus. Glass coverslips are coated with L-lysine overnight at 22°C under UV light in a 12 well plate. Post coating HUH7 cells are seeded onto the coverslips and grown till 50 % confluent. The cells are transfected with M1wt, M1K21R, and M1K242R. Post 24 hours of transfection, the cells are washed with DPBS, and fixed in 4% Paraformaldehyde (PFA) for 15 minutes with rocking. Post fixing the PFA is aspirated, and the cells are washed with DPBS. After fixing the cells are blocked (1XDPBS, 1 % BSA, 0.1 % Triton x-100) for 60 minutes at 22°C with rocking. Post blocking the antibody is diluted 1:100 in blocking buffer and is stained overnight at 4°C with rocking. The cells are rinsed with DPBS for 5 minutes and repeated 3 times. The cells are then incubated for 60 minutes with the secondary anti-mouse 488 Alexa-dye (Invitrogen) in 1XDPBS, 1 % BSA, 0.1 % Triton x-100. The cells are rinsed with DPBS for 5 minutes and repeated 3 times. Post-secondary stain the cell nucleus was stained with Hoechst 33342 for incubation of 15 minutes. Post nuclear stain, the cells are washed 4 times with DPBS with 5-minute incubation. The cells were imaged on Olympus BX43, and images were stacked and analyzed using ImageJ software.

### qFRET K_D_ of SUMOylated N Protein

The evaluation of N protein oligomerization by *in vitro* qFRET based K_D_ affinity assay. The individual N protein wild type and mutants were first cloned into the FRET fusion genes. Each N protein was tagged with the donor or acceptor pair fluorescent proteins for the implementation of the qFRET assay. Thus, five pairs of N protein genes, the wild type, Lys mutations on individual 61, 65, 347, and 355, were cloned into pET-28(b) containing CyPet or YPet, respectively. Each pair of proteins were SUMOylated *in vitro* following optimized concentrations of SUMO enzymes and SUMO1 as following, 6xHisCyPet-SUMO1 6 *μ*M, 6xHisYPet-N protein 6 *μ*M, E1 hetro-dimer AOS1/UBA2 at 100 nM, E2 conjugating enzyme UBC9 200 nM, E3 ligase PIAS1 250 nM in a SUMOylation buffer (20 mM Tris-HCl (pH 7.5) 50 mM NaCl, 4 mM MgCl, 1 mM DTT). Negative control, without ATP, was set up for non-specific interaction. The SUMOylation reaction was incubated at 37°C for 1 hour, along with the negative control of reaction without ATP.

The reaction included the N protein bound to CyPet (donor) and the separate reaction of bound YPet (acceptor) after the SUMOylation. The SUMOylated N proteins were directly measured for affinities. The qFRET-based K_D_ method contained the donor fusion protein concentration at 500 nM, and the acceptor is titrated from 0 nM to 2,500 nM. The series of titrations were individually performed to determine the E_mFRET_ values of each YPet-N protein. Following Equation 1 we measured the three wavelengths, E_mTotal_, FL_DD_, and FL_AA_ for each reaction. The measurements of were measured on Molecular Devices Spectra M3™, with “Endpoint” settings and PMT at constant gain set to “Low”. The K_D_ was then determined using the multi-regression method in Equation 2:

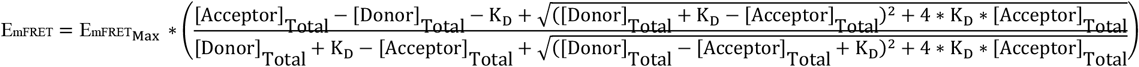

For the data analysis, GraphpadPrism5™ was used to fit the nonlinear equation to the data set collected. The constraints for the non-linear regression fit were set to donor concentration at a constant of 500 nM, and the constraint K_D_ and E_mFRETMax_ cannot be zero.

## Results

### *In vitro* qFRET Assay for SUMOylation of N Protein

We have developed an in vitro qFRET-based SUMOylation assay including SUMO E1 activating enzyme, E2 conjugation enzyme, and E3 ligase. In this assay, the SUMO peptide was fused with the FRET donor, CyPet, and the substrate SARS-CoV-2 was fused with the FRET acceptor, YPet (Fig.1A). In the presence of SUMOylation enzymes, the CyPet- SUMO1 is conjugated to the YPet-N protein, leading to a FRET signal (Fig.1B). We validated this assay using a classical Western-blot assay(Fig.1C). The SUMOylation of N protein was significantly enhanced in the presence of E3 ligase, PIAS1, indicating by the disapprance of free CyPet-SUMO1(Fig. 1C). The results suggest that the FRET-based SUMOylation assay is feasible and valid.

**Figure 1.**
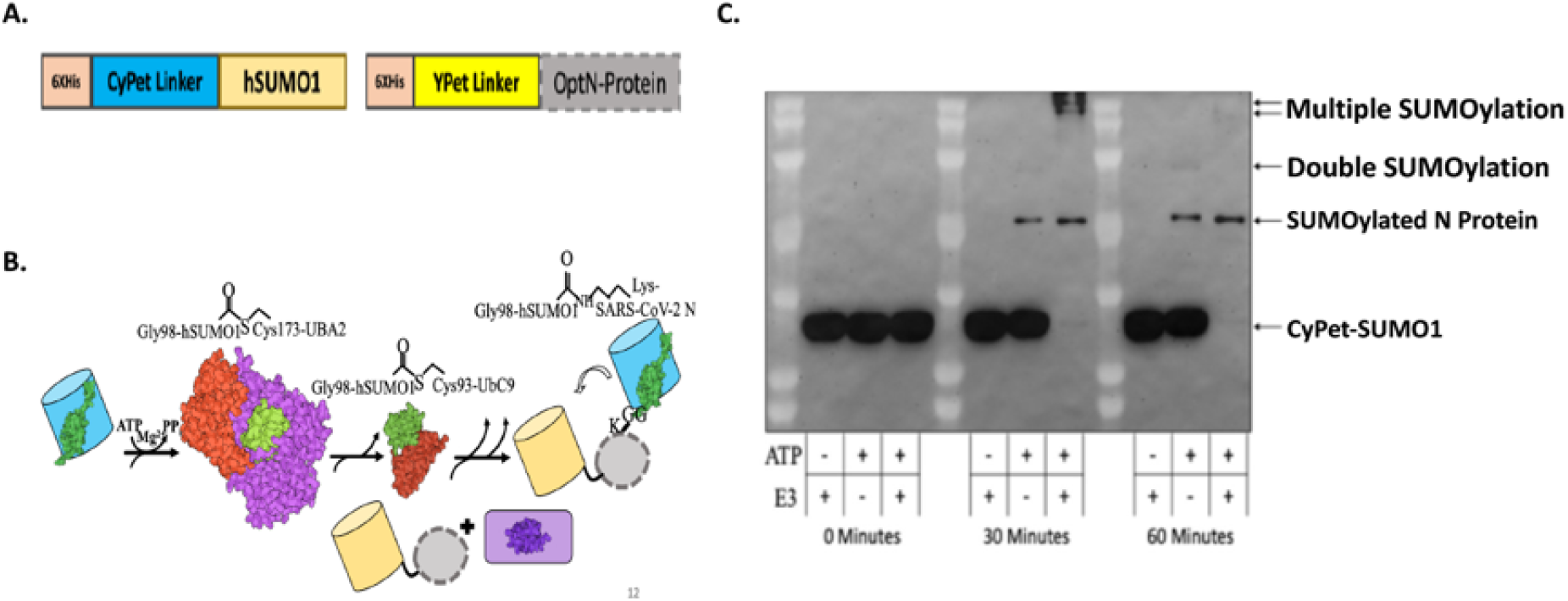
SUMOylation of SARS-CoV-2 N protein. A. Diagram of the fluorescence fusion protein CyPet-SUMO1 and YPet-N for the qFRET-based SUMOylation assay. B. *In-vitro* SUMOylation assay with the FRET as a reporter signal. The fusion protein CyPet-SUMO is first bound to E1 activating enzyme, for the intermediate E1-Cypet-SUMO1 thioester bond at cys-173 to glycine 98 on SUMO1. The SUMO is then transferred to the catalytic cystiene93 of E2 conjugating enzyme. The E3 ligase and target protein are said to non-covalently interact with the E2-SUMO1 complex. The CyPet-SUMO1 is then shuttled to a lysine on the target protein, to be covalently bound by an isopeptide bond. C. The Western-blot of N protein SUMOylation in the in vitro reaction containing E1, E2 and E3.

### Mass Spectrometry Analysis to Determine SUMO Modified Lysine on N Protein

We then conducted the SUMOylaiton assay of SARS-CoV-2 protein using qFRET to follow the SUMO peptide conjugation. The observed E_mFRET_ signal for the SUMOylation of SARS-CoV-2 N protein showed a significant difference in the magnitude of E_mFRET_ without or with the addition of ATP (Fig.2A). A t-test analysis was completed on the triplicate measurements, compared with and without ATP, as well as with and without E3 ligase. There was also a significant E_mFRET_ signal increase in the presence of E3 ligase, suggesting a robust SUMOylation event. A one-way ANOVA was used to analyze the significance between the control group without ATP (-ATP), with ATP and no E3 ligase, and with ATP and E3 ligase.

### We then determined the SUMOylation sites of SARS-CoV-2 N protein using

Thermo Orbitrap Fusion. The identified modified lysines are illustrated in Fig.2B along with the other 25 lysines found unmodified by the SUMO1. The coverage of the N protein was 95 %. The spectrum of the identified peptide with lysine 61 modification was a large section of N protein from position 41 to 67, which had Lys 61 and 65 contained within it (Fig.2C). This cut pattern matched for n-terminal arginine-41, and the cleave at arginine-68 after the proline-67. The GluC protease cutted the carboxyl ends of glutamic acids and could also cut the n terminal of arginine (R) and c terminal of aspartic acid (D). The SUMO1 peptide “GGTQ” followed by a glutamic acid at position 61 was cleaved and matches the expected GluC cut. Fig. 2D has a similar peptide however with lysine-65 holding the SUMO1 modification. The GluC cut positions on the peptide were the same as the previous peptide, N protein position 41 - 67. The N protein peptide from positions 341 – 355 showed SUMO1 GG peptide modifications (Fig.2E). This peptide matched the GluC cut pattern at aspartic acid position 341 – 357. However, some discrepancies were noted in, the c terminal of the peptide should have an isoleucine after the modified lysine. This could be an aberration in the MS analysis or an artifact of GluC non-specific cut. We still included these two sites in the study as the other 25 lysines had no detected modification.

**Figure 2.**
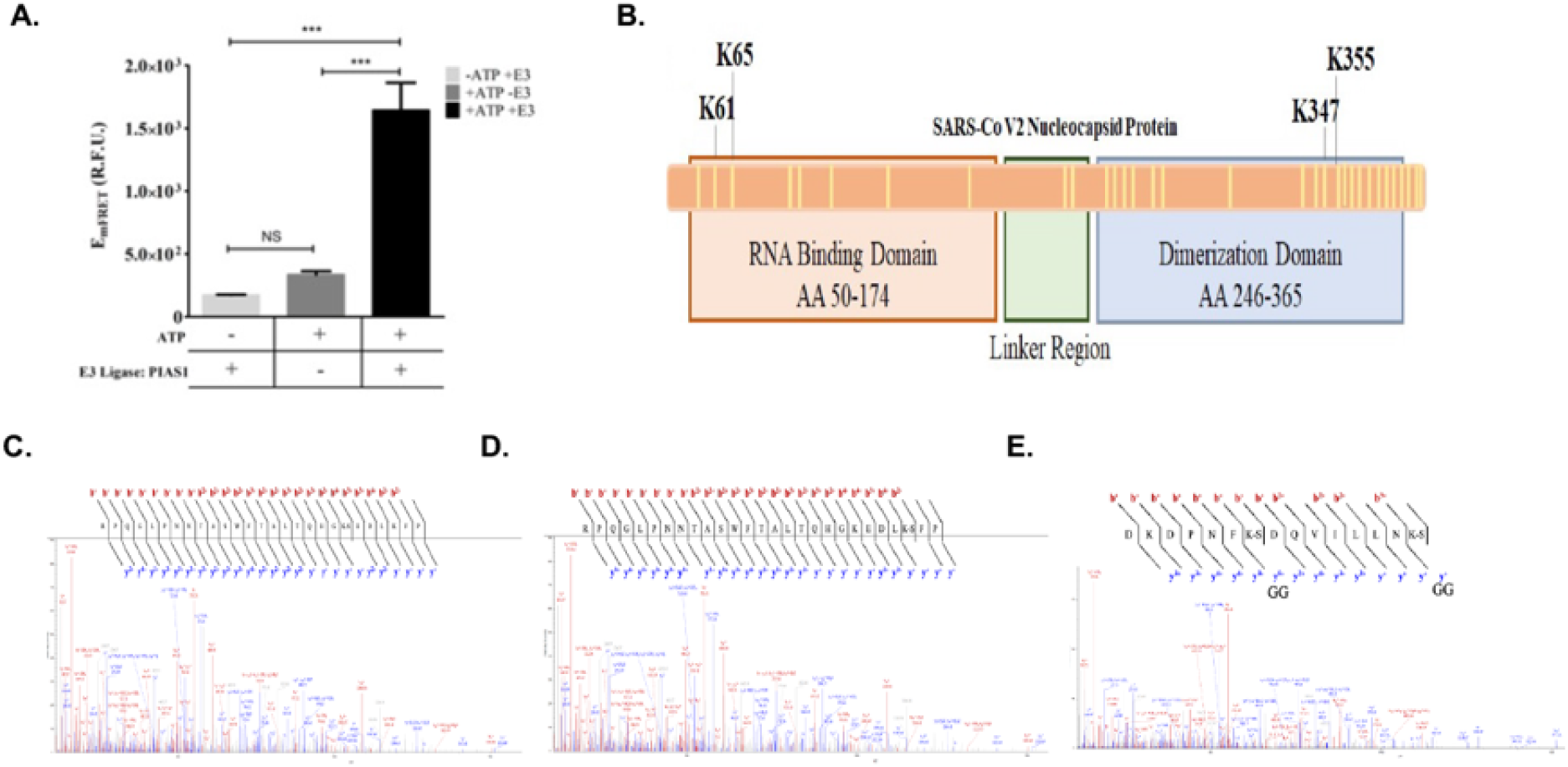
*In vitro* SUMOylation of SARS-CoV-2 N protein and SUMOyaltion site identification using Mass Spectrometry(MS). A. The *in vitro* MS sample was measured for qFRET signal before processing for MS, with and without ATP or E3 PIAS1. B. The illustration of the location of the four discovered lysines, along with a total of 31 lysines on the protein shown as yellow lines. C. The spectrum were generated by Thermofisher Proteome Discoverer™ MS spectrum of peptide containing modified K61. D. The spectrum of K65 peptide with GluC cut on SUMO1, GGTQ. E. The spectrum of both K347 and K355 found in the same peptide, with SUMO1 peptide GG cut with GluC.

The secondary analysis of each lysine site can be done through evaluation of each site matching the SUMO consensus motif. Based on the SUMO consensus motif, of a hydrophobic residue (Ψ), the modified lysine (K), any amino acid, and either an aspartic acid or a glutamic acid. The Ψ-K-x-D/E motif has been applied by numerous groups to ascertain SUMOylation site. Using two only servers, GPS-SUMO and JASSA were used to determine if these sites matched the SUMO consensus(19, 20). Both servers only pointed to lysine position 338 as the highest probability, and lysine 61, 65, 347, and 355 all were low probability. This result did not match the SARS-CoV-1 N protein SUMOylation site, also does not match the in vitro SUMOylation sites, and was not included in this study.

### qFRET Assay for *In-Vitro* SUMOylation N Protein Mutants

The results of the *in vitro* SUMOylation of N protein with qFRET as a reporter was then performed in order to show the SUMOylatio site modifications (Fig. 3). We observed significant signal from wildtype N protein with E3 ligase and ATP, indicating a good activity of PIAS1 ligase, for N protein (Fig.3A). In comparison the wildtype N protein, the 61 and 65 mutants showed a significant drop in the signal without E3, and this pattern was same as in the double K61 and K65mutant without E3 ligase. However, both K61R and K65R still showed significant FRET signal, indicating SUMOylation occurred at other Lys sites. Both K347R and K355R mutants of N protein still showed significant FRET signals, suggesting other Lys sites, such as K61 or K65, were still SUMOylated. The significant differences could be seen in reactions without E3 ligase or without ATP. The one-way ANOVA analysis with Tukey Test, results in significant differences when E3 was added across all reactions.

**Figure 3.**
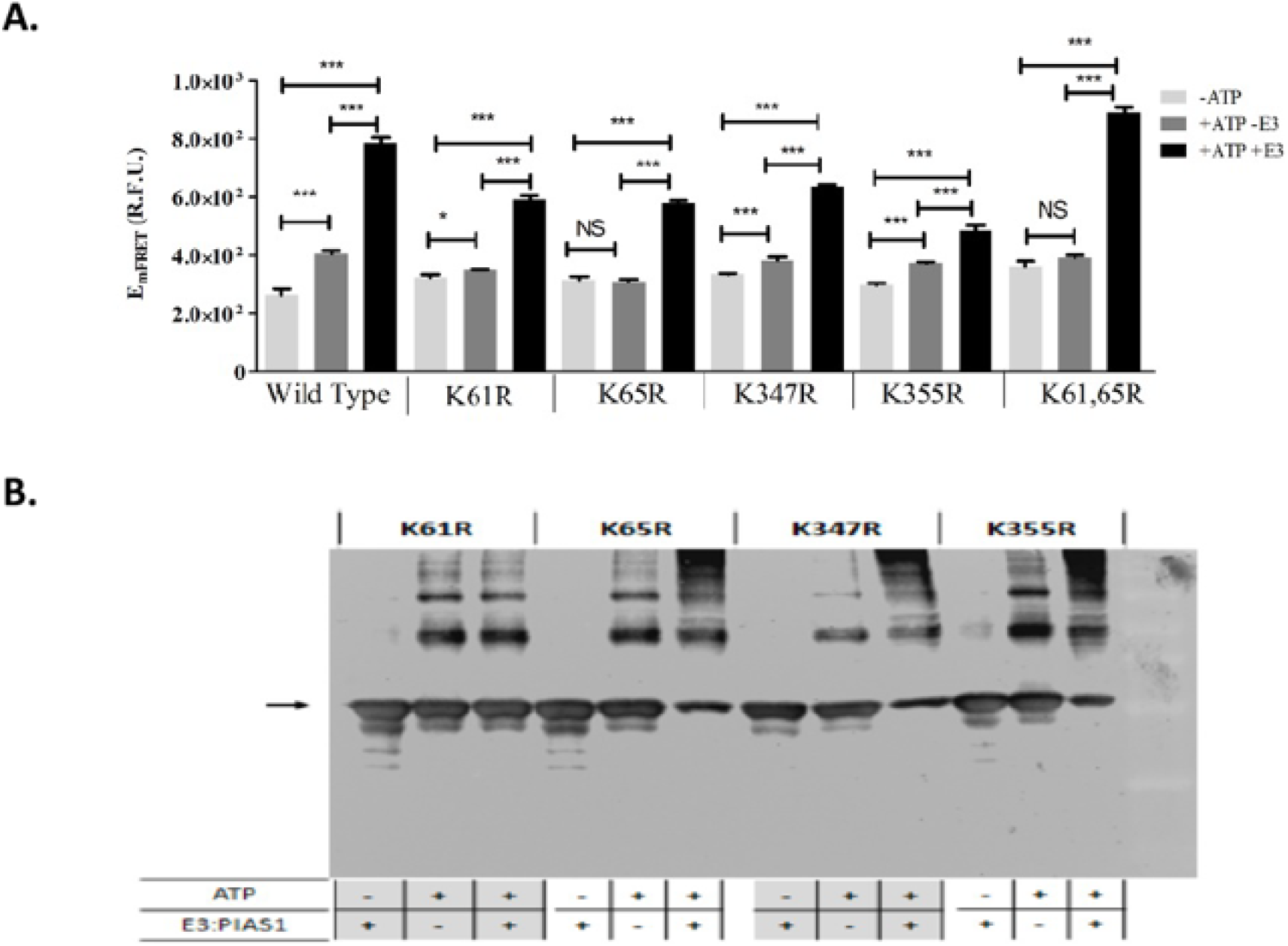
In vitro SUMOylation of N protein and its Lys mutants. **A**. SUMOyaltion assay of wildtype and K mutants of N protein using qFRET assay. Comparison of no ATP (-ATP), with no E3 ligase PIAS1 (-E3), and a complete reaction with ATP and E3 ligase PIAS1 (+ ATP+ E3). The reactions were all done under the same conditions, and the measurements were all taken on the same instrument, Molecular Devices SpectraMax3™. One-Way ANOVA was done on the data sets of -ATP/-E3/+ATP+E3, the -ATP was the control group. Tukey test was used as the post-HOC analysis. P values are p<0.0001***, p<0.05*, and no significant difference (ns), n=3. **B**. The Western-blot of the wildtype and K mutants of N protein.

### qFRET K_D_ of SUMOylated N Protein

Because the SARS-CoV-2 protein interact with each other to form oligomers for viral genome RNA packaging, the interaction of N protein with each other is critical for its functions *in vivo*. We therefor performed qFRET assays to determine the interaction affinities of N wildtype and Lys mutant proteins with or without SUMOylation. One of the advantages of our qFRET technology does not require very pure proteins for the K_D_ determination. Therefore, we conducted the *in vitro* SUMOyaltion of all N proteins with CyPet-SUMO1 and YPet-N proteins first and followed with the K_D_ determinations without purification(Xiong et.al unpublished).

The E_mFRET_ response from the titration of total acceptor protein and the fit profile plotted in Fig.4. The E_mFRET_ of wildtype N protein from the titration of acceptor fusion protein YPet-Nwt was plotted in points (circle/orange) for unmodified, and (diamond/green) for SUMO1 modified (Fig.4A). We observed a K_D_ value of wildtype N protein interaction with itself after SUMO1 modification to be 0.46 µM and standard error of 0.34, and 1.46 µM standard error of 0.09 when not modified(Fig.4B). This result suggests that SUMOylaiton modification of SARS-CoV-2 N protein increases its interaction affinity with itself, and this modification may help the oligomerization of N protein in nucleus for viral RNA genome packaging.

**Figure 4.**
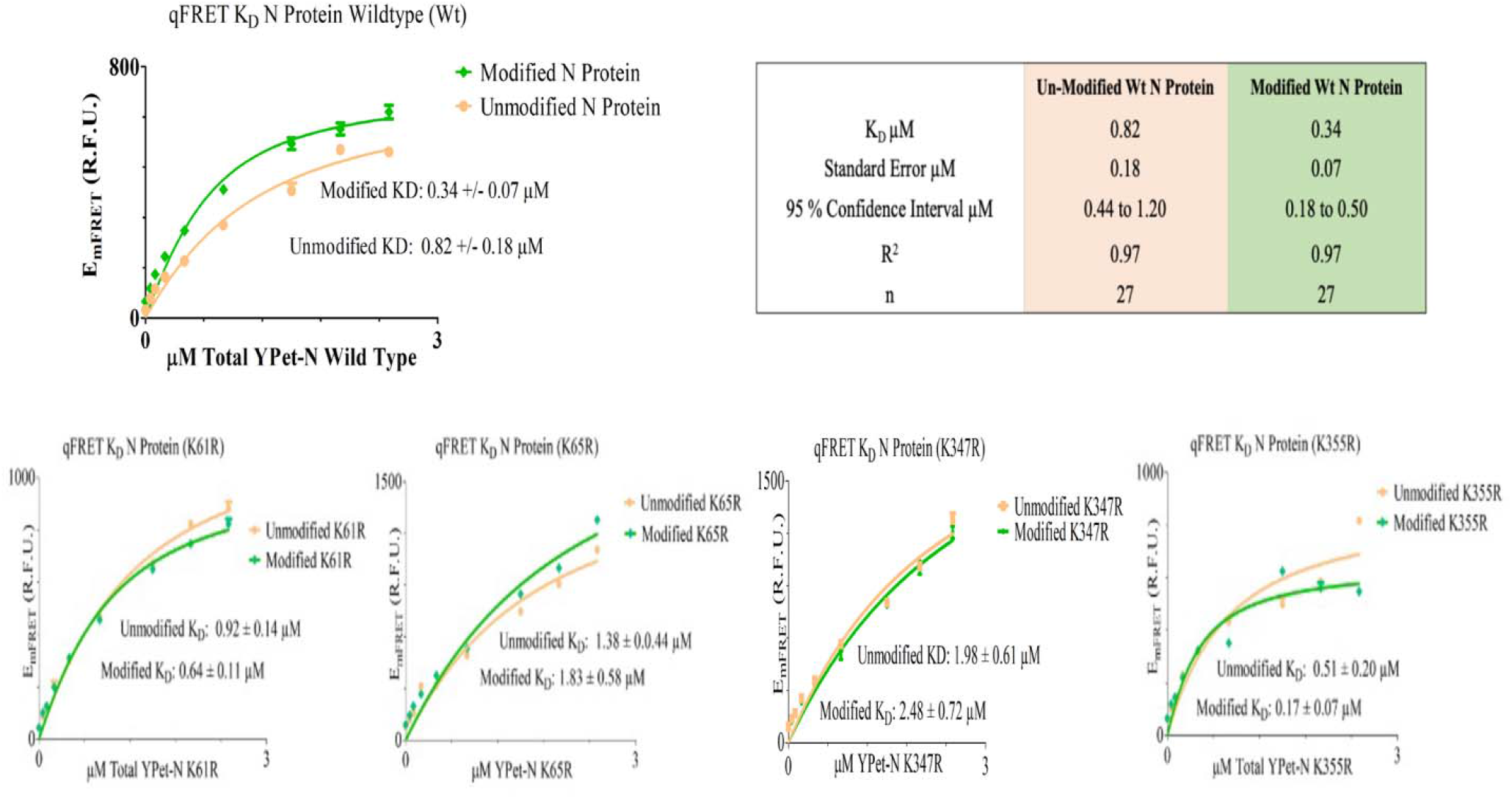
The K_D_ determinations of SUMOylated or un-SUMOylated N protein-N protein interactions using qFRET assay. **A**. The binding curve of wildtype of N protein with or without SUMOylation. The EmFRET signal fit of SUMO modified(Diamond/Green), and unmodified (Circle/Orange). **B**. The Binding curves of the SUMO modified N protein mutants of (K61R). C. The Binding curves of the SUMO modified N protein mutants of (K61R). **D**. The Binding curves of the SUMO modified N protein mutants of (K65R). **E**. The Binding curves of the SUMO modified N protein mutants of (K347R). **F**. The Binding curves of the SUMO modified N protein mutants of (K355R). (Diamond/Green) and the unmodified (Circle/Orange). All plots and the fits were generated on GraphpadPrism5™.

We then tried to determine which Lys residure SUMOylation contribute to the interaction affinity increase. The N protein mutants at each Lys residue position, 61,65, 347 and 355, was mutated to Arg, then were expressed and SUMOylated before the K_D_ determination. The results of qFRET K_D_ assay on the N protein mutants are listed in Table 4. Both modified and unmodified N mutant protein were determined for K_D_ values with or without SUMOylation modification. The E_mFRET_ values across 0 to 2.5 µM of YPet-N protein were determined with and without SUMOylation. The affinity of the un-modified N proteins ranged from 0.65 to 2.45 µM, and the range for modified was 0.30 to 1.70 µM for all four Lys residues. Although the interaction affinities of unmodified N proteins are slightly different may be due to some conformational changes after Lys residue mutations, the results show a general increase of interaction affinity with SUMOylation modification. The combination of SUMOylation of three Lys residure, 61,65 and 347 seems contribute most to the affinity increase.

**Table 4.** The K_D_ values of the single mutant SARS-CoV-2 N protein with and without SUMOylation modification.

| N protein | K61R<br>Unmodified | K61R<br>Modified | K65R<br>Unmodified | K65R<br>Modified | K347R<br>Unmodified | K347R<br>Modified | K355R<br>Unmodified | K355R<br>Modified |
| --- | --- | --- | --- | --- | --- | --- | --- | --- |
| $K_D$ (μM) | 0.92 | 0.64 | 1.38 | 1.84 | 1.98 | 1.70 | 0.51 | 0.17 |
| Standard Error (μM) | 0.14 | 0.11 | 0.44 | 0.58 | 0.61 | 0.37 | 0.20 | 0.07 |
| 95 % Confidence Interval (μM) | 0.62 to 1.21 | 0.41 to 0.87 | 0.46 to 2.30 | 0.64 to 3.03 | 0.71 to 3.21 | 0.99 to 3.97 | 0.08 to 0.90 | 0.01 to 0.32 |
| $R^2$ | 0.99 | 0.98 | 0.95 | 0.91 | 0.97 | 0.97 | 0.90 | 0.90 |
| n | 27 | 27 | 27 | 27 | 27 | 27 | 27 | 27 |

### Nucleus Translocation of N Protein

We then examined the potential role of SUMOylation on each Lys site for its nucleus translocation of the SARS-CoV-2 N protein. We used the immunofluorescence protein YPet as imaging marker of fusion protein YPet-N to tract the sub-cellular localization of wildtype or Lys mutant N protein. The images at the top row were the nuclear stains using Hoechst that provide the location of the nucleus(Fig.5). The YPet-N fluorescence image showed universal distributions among the whole cells including both cytosol and nucleus, but with a very obvious condensated granules. The images at the second row were obtained through the YPet channel, where we observe the YPet fused N protein. The formed bright spots were observed to be condensate or granules of N protein. These fluorescent granules forming within the cell are inherent to the N protein function as oligomers. The first column of images were images of the wild type N protein, and we observe granules/condensated particles in the nucleus and the cytosol. The N proteins with three Lys mutations, K61R, K347 and K355, were all observed as granules in both cytosol and nucleus. Interstingly, the N K65R protein showed predominantly in the cytosol, not in the nucleus, suggesting that Lys residue 65 SUMOylation may be essential for its nucleus localization. This result indicates that SUMOylation of N protein may play a critical for its nucleus translocation and subsequently viral RNA genome packaging.

**Figure 5.**
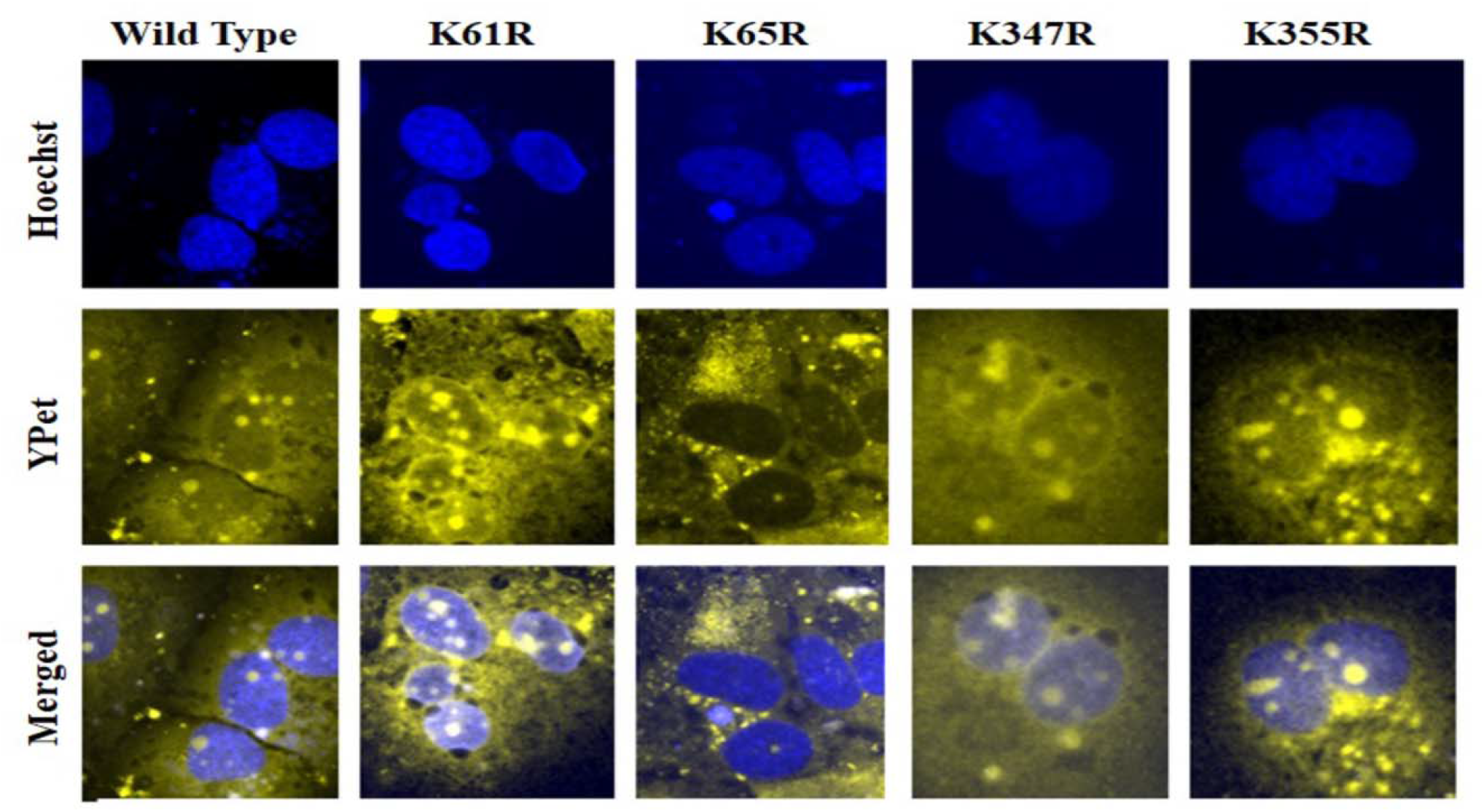
Subcellular localization determination of wildtype N protein and Lys mutants using a fluorescent microscope. The nuclear stain Hoechst was determined at 488 nm. The images of YPet-tagged-N proteins and its mutants were taken at at 533 nm using the fluorescent microscope, Olympus BX43.. Images were processed on ImageJ™.

## Discussion

It has been generally observed that SUMOylation has been extensively involved in the host-virus interactions and viruses manipulate host SUMOylaiton for their benefits of relications in host cells(21-24). We find that SUMO modification can significantly enhance its interaction affinity with itself. The N protein is found to crosslink at high concentrations in the cell, but forms tetramers and dimers(9, 10). The overall organization of the vRNP complex formation by the N protein requires non-covalent interaction of the various forms of N protein. Demonstrated here that the modulation of affinity of N protein with the SUMO modification may contribute significantly to its oligomerization. The qFRET platform can provide a rapid assessment of a protein interaction affinity of viral proteins. It will be very interesting to see whether the SUMOylation inhibitor can interfere with the SARS-CoV-2 replication.

We identified four SUMOylation sites, K61, 65, 347, and 355 of SARS-CoV-2 Nucleocapsid protein using in *in vitro* SUMOylation assay SUMOylation assay in the presence of SUMO E1, E2 and E3 enzymes. The qFRET-based SUMOylation assay, used alternatively but more sensitive and robustness to immunoblot, is used in the evaluation of covalent modifications of proteins. The image of fluorescence protein tagged N protein forms bright spots as granules in cells. We observe the wildtype N protein in both the cytosol and the nucleus, consistent with previous report of SARS N protein. These findings are consistent with reports of N protein cellular translocation. The alteration of nuclear translocation of N protein was observed with lysine 65 mutation, which mainly restricts protein in the cytosol. The finding suggest a decrease of N protein nuclear translocation without SUMOylation may impact the viral genome packaging in the nucleus.

The SARS-CoV-2 proteome is relatively new to the scientific community, and access to versatile techniques that can evaluate protein modifications and evaluate protein properties is difficult. Demonstrated here is a versatile method to evaluate the covalent modification and non-covalent interactions using the same platform. An advantage of the *in vitro* evaluation of SUMOylation modifications is it also provides certainty in the observations. Due to the numerous in-cell lysine modifications the probability of missing or false negative classification of a lysine modification is high. Furthermore, the yield of a modified protein from an in-cell pull-down assay can be challenging and have low yields. Thus, researchers look to over express the SUMOylation mechanism, however within a cellular environment, over SUMOylation can bring about unwanted consequences to the cellular proteome. The *in vitro* qFRET assay used here provided a fluorescent reporter for SUMOylation and was directly used in the identification of modified lysine residues. The coverage of the protein identified in MS was up to 95 %, the other 5 % is assumed to be degraded during the digestion and sample preparation. The high certainty and the overall coverage of the protein in MS analysis provide confidence in the identified SUMOylation sites.

The reconstitution of SUMOylation reaction with qFRET as a reporter is a robust and rapid method for identifying SUMOylation events, characterizing enzymatic activity, and identifying SUMOylation sites(18, 25, 26). This method includes the E3 ligase PIAS1 for its enhanced SUMOylation activity and is coupled with mass spectrometry to provide insight into identifying multiple SUMOylation sites(27-29). The qFRET assay utilizes a FRET optimized donor fluorescent protein tag, CyPet, on the SUMO1 protein and FRET optimized acceptor fluorescent protein tag, YPet, on the N-protein(30). The CyPet-YPet fluorescent proteins experience the non-radiative FRET phenomenon, when within 10 nm distance between them which is applicable for observing SUMO1 attachment. The FRET phenomenon has been described and applied in various studies of protein-protein interaction, and especially to measure molecule distance as FRET efficiency is proportional to the r^6^ between the two fluorophores(31-33). We have developed a fluorescence-based method in determining covalent attachment of Cypet-SUMO1 to YPet-N protein. The method demonstrated here applies a “three cube FRET” fluorescence reporter that extracts the emission of FRET signal, E_mFRET_, from the raw fluorescent signal, EmTotal, at the FRET wavelength. The extraction applies three different fluorescent measurements (Table 1), to extract the E_mFRET_ response from a FRET reaction. The method filters out cross channel signal of the unbound donor or acceptor from the FRET wavelength. The relationship applied in Equation 1, determines the contribution of cross talk form both acceptor and donor by applying ratiometric constants alpha (α Equation 2) and beta (β Equation 3), to subtract the crosstalk signal from the EmTotal. The details on the development of the qFRET method can be found on a previous study on development of the qFRET signal by Song et al. (2011)(18). The method has been applied previously to determine kinetic values of protein-protein interactions such as dissociation constant K_D_, enzymatic constants k_cat_/K_m_, and applied to assess SUMO modification of viral proteins(18, 26, 33).

## Conclusions

Viruses take advantages of host factors for their infection, replication, virion assembly and budding for their amplification and consequently pathogenesis, and therefore targeting host factors as a new strategy for anti-virus therapeutics is very promising(34-38). Human SUMOylation has been extensively utilized by various viruses, including influenza A/B virus, HIV and Ebola viruses(22, 23, 39). Here, we identified the SUMOylation sites of SARS-CoV-2 N protein, which is critical for viral genome RNA packing, using an *in vitro* qFRET-MS coupled approach. Among these SUMOyaltion sites, the K65R mutation prevents the nucleus translocation of N protein. The K_D_ values of interactions of wild type and Lys mutants of N proteins have also been determined using the qFRET assay in solution. It shows that the SUMOylation can significantly increase the integration of N protein with itself. This novel discovery supports the roles of N protein in viral RNA genome packaging as oligomers. These discoveries could provide new insights for human-SARS-CoV-2 interactions, which can be potential a novel strategy for inhibiting host factors as anti-viral therapeutics development.

## Author Contributions

Conceptualization, J.L.; methodology, J.L., V.G.J.R., V.M.; validation, J.L.,V.M.; formal analysis, V.M., J.L.; investigation, V.M.; resources, J.L.; data curation, V.M., J.L.; writing—original draft preparation, V.M., J.L.; writing—review and editing, J.L.; visualization, V.M.; supervision, J.L.,V.G.J.R.; funding acquisition, J.L. All authors have read and agreed to the published version of the manuscript.

## Funding

This work was partially funded by the UCR Academic Senate Research Grant and Attaisina Gift grant to JL.

## Data Availability Statement

All the data can be available upon request to J.L. at.

## Acknowledgments

We appreciate Zhehao Xiong and George Way for their pioneer work in SUMOylation inhibitor for IAV and IBV virus drug.

## Conflicts of Interest

The J.L. laboratory has received research support from Attaisina. J.L. and V.M. are inventors on patents and patent applications on the use of SUMOylation inhibitor for virus infections and cancer, owned by the University of California at Riverside, outside of the reported work. The funders had no role in the design of the study; in the collection, analyses, or interpretation of data; in the writing of the manuscript, or in the decision to publish the results.

